# A positive correlation between GC content and growth temperature in prokaryotes

**DOI:** 10.1101/2021.04.27.441598

**Authors:** En-Ze Hu, Xin-Ran Lan, Zhi-Ling Liu, Jie Gao, Deng-Ke Niu

## Abstract

**Background:** GC pairs are generally more stable than AT pairs; GC-rich genomes were proposed to be more adapted to high temperatures than AT-rich genomes. Previous studies consistently showed positive correlations between growth temperature and the GC contents of structural RNA genes. However, for the whole genome sequences and the silent sites of the codons in protein-coding genes, the relationship between GC content and growth temperature is in a long-lasting debate.

**Results:** With a dataset much larger than previous studies (681 bacteria and 155 archaea with completely assembled genomes), our phylogenetic comparative analyses showed positive correlations between optimal growth temperature (Topt) and GC content both in bacterial and archaeal structural RNA genes and in bacterial whole genome sequences, chromosomal sequences, plasmid sequences, core genes, and accessory genes. However, in the 155 archaea, we did not observe a significant positive correlation of Topt with whole-genome GC content (GC_w_) or GC content at four-fold degenerate sites. We randomly drew 155 samples from the 681 bacteria for 1000 rounds. In most cases (> 95%), the positive correlations between Topt and genomic GC contents became statistically nonsignificant (*P* > 0.05). This result suggested that the small sample sizes might account for the lack of positive correlations between growth temperature and genomic GC content in the 155 archaea and the bacterial samples of previous studies. Comparing the GC content among four categories (psychrophiles/psychrotrophiles, mesophiles, thermophiles, and hyperthermophiles) also revealed a positive correlation between GC_w_ and growth temperature in bacteria. By including the GC_w_ of incompletely assembled genomes, we expanded the sample size of archaea to 303. Positive correlations between GC_w_ and Topt appear especially after excluding the halophilic archaea whose GC contents might be are strongly shaped by intense UV radiation.

**Conclusions:** This study explains the previous contradictory observations and ends a long debate. Prokaryotes growing in high temperatures have higher GC contents. Thermal adaptation is one possible explanation for the positive association. Meanwhile, we propose that the elevated efficiency of DNA repair in response to heat mutagenesis might have the by-product of increasing GC content like that happens in intracellular symbionts and marine bacterioplankton.

## Background

As guanine (G) strictly pairs with cytosine (C) and adenine (A) pairs with thymine (T) in DNA double helix, the amount of G is equal to C, and that of A is equal to T in the genomes of any cellular organisms. GC content, i.e., the percentage of G + C, is widely used to measure genomic nucleotide composition. It is a highly variable trait ranging from 8% to 75% (1–3). This genomic trait has been widely studied, and its evolution has been proposed to be associated with numerous mutational and selective forces driven by genetic, metabolic, and ecological factors (4–19). The high temperature might be the most long-debating (20–22). Because G:C pairs have an additional hydrogen bond than A:T pairs, the GC-rich genomes are expected to be thermally more stable in high-temperature environments (23). Bernardi and Bernardi (24) proposed that high GC content is a thermal adaptation of warm□blooded animals.

As prokaryotes have a much wider thermal distribution than plants and animals, bacterial and archaeal genomes are the best materials to test the thermal adaptation hypothesis. An analysis of 764 prokaryotic species, including mesophilic genera and thermophilic genera, did not find a correlation between whole-genome GC content (GC_w_) and the optimal growth temperature (Topt) (22). However, this study found a significant positive correlation between Topt and the GC content of structural RNAs (tRNAs and rRNAs). The rationale of these observations is that the secondary structures of tRNAs and rRNAs are more sensitive to high temperatures than the double-strand helix of DNA. In most prokaryotes, protein-coding genes take most of the genome size. Protein structures and functions constrain the GC content evolution at the nonsynonymous sites of the codons. This functional constraint might conceal the hypothetical thermal adaptation. Compared with GC_w_, the GC content at the third site of the codons (GC_3_) is more desirable to test the thermal adaptation hypothesis. Early solitary cases indicated that GC_3_ might be related to growth temperature. For example, the tyrosy1□tRNA synthetase gene isolated from the thermophile *Bacillus stearothermophilus* (current name: *Geobacillus stearothermophilus*) has a higher GC_3_ than the homologous gene in *Escherichia coli*, 68.0% *vs.* 59.4% (25). The *leuB* gene isolated from the extreme thermophile *Thermus thermophilus* HB8 has an extremely high GC_3_, 89.4% (26). Hurst and Merchant examined the relationship between GC_3_ and Topt of 29 archaeal species and 72 bacterial species for a general conclusion (27). They did not find significant correlations between Topt and GC_3_ or Topt and GC_w_. At the same time, they also found a significant positive correlation between the GC content of structural RNAs and the Topt in both archaea and bacteria. Their analysis accounted for the effect of shared ancestry, so they provided more robust evidence against the thermal adaptation hypothesis. Soon afterward, Xia et al. (28) showed that the growth at increasing temperature (from 37°C to 45°C) for 14,400 generations did not increase but decreased the genomic GC content of the bacterium *Pasteurella multocida*. Furthermore, Lambros et al. (29) reported a negative correlation between optimal growth temperature and the GC content of protein-coding genes in 550 prokaryotes. As the effect of the shared ancestry had not been controlled in their study, we must be cautious in response to their results because the potential nonindependence among their data might violate the basic assumption of the statistical models used in their study.

Subsequently, Musto et al. (30) published a debate-provoking study. As many environmental factors likely influence genomic GC content evolution, closely related species are expected to differ in fewer environmental factors than distantly related species. The correlation of GC content with growth temperature is less likely disturbed by other factors when the analysis is limited within closely related species. Therefore, Musto et al. (30) examined the relationship between genomic GC content and Topt with each prokaryotic family. Among the 20 families they studied, the number of families with positive correlations is significantly higher than expected by chance, no matter the effect of the common ancestors was accounted for or not. Meanwhile, they observed a significant positive correlation when considering all independent contrasts from different families together. However, Marashi and Ghalanbor (31) noticed that most of the significant correlations within each family depend heavily on the presence of a few outlier species. Exclusion of only one species would lead to loss of significant correlations in several families. Basak et al. (32) pointed out that the correlation is sensitive to the presence or absence of a few outliers in some families because the sample sizes in these families were too small. Using non-parametric correlation analysis that is not sensitive to the presence of outliers, Musto et al. (33) repeated their analysis and confirmed their previous results. The debate did not end after that. Wang et al. (34) updated the Topt values for some species and found that the positive correlation between Topt and genomic GC content in two families disappeared. Besides, they suggested that the positive correlation between Topt and genomic GC content in the family Enterobacteriaceae should be explained by the correlation between genome size and optimal growth temperature. Still, this study did not shake the confidence of Musto et al. (35) on the correlation between Topt and genomic GC content in prokaryotes. Although Musto and coauthors have rebutted all the criticisms, their studies have not convinced later authors of review articles (4, 20, 36). For example, Agashe and Shankar (4) claimed that “*it seems unlikely that genomic GC content is driven by thermal adaptation*” after reviewing the results of Hurst and Merchant (27) and Xia et al. (28), but without mentioning the debates on Musto et al. (30).

As prokaryotic genomes often have many accessory genes frequently lost and gained, the genome-wide measures of GC content could roughly reflect the shaping effects of environmental factors in evolution. By contrast, the structural RNA genes ubiquitously exist in prokaryotic genomes, and their GC contents are more comparable in large-scale phylogenetic analyses. Similarly, the core genome or strictly defined orthologous genes could also accurately reflect the historical shaping effect of growth temperature on GC content evolution. Ream et al. (37) analyzed the GC contents of two genes (*ldh-a* and *α-actin*) across 51 vertebrate species with adaptation temperatures ranging from −1.86°C to approximately 45°C. They did not find any significant positive correlations between living temperature and GC content, whether the GC content is measured by the entire sequences, the third codon position, or the fourfold degenerate sites. However, Zheng and Wu (38) found a positive correlation between growth temperature and the GC content in the coding regions of four genes across 815 prokaryotic species, including mesophiles, thermophiles, and hyperthermophiles. These four genes shared by all the 815 prokaryotic genomes could be considered strictly defined core genomes.

Using a manually collected dataset of growth temperature and without accounting for the effect of the common ancestors, Sato et al. (39) recently confirmed the results of Galtier and Lobry (22). It should be noted that the correlation between Topt and the GC content of structural RNA was consistently observed in much more studies than those mentioned above (19, 39–43). By contrast, as reviewed above, the correlation between Topt and genomic GC content, if it exists, depends heavily on the sample size, the families of prokaryotes, the sequences, and the methods used to detect it.

Benefitting from the manually curated growth temperature dataset from the database TEMPURA (39), we comprehensively analyzed the relationship between growth temperature and GC content. The present study covers three indexes of growth temperature (maximal growth temperature [Tmax], Topt, and minimal growth temperature [Tmin]) and a series of GC content indexes, including GC_w_, GC content of the protein-coding sequences (GC_p_), GC content at fourfold degenerate sites (GC_4_), GC content of the genes coding structural RNAs (tRNA, GC_tRNA_; 5S rRNA, GC_5S_; 16S rRNA, GC_16S_; 23S rRNA, GC_23S_) and GC content of non-coding DNA (GC_non_, including intergenic sequences and untranslated regions of mRNA that are generally unannotated in prokaryotic genomes). The whole genome, primary chromosome sequences, plasmid sequences, core genes, and accessory genes have been examined separately. Our results support a positive correlation between genomic GC content and growth temperature in bacteria and likely in archaea.

## Results

### Strong phylogenetic signals in both GC contents and growth temperatures

A significant force shaping prokaryotic evolution is horizontal gene transfer, making the genealogical relationships among bacteria and archaea exhibit a somewhat network-like structure. If bifurcation is not the phylogeny’s dominant pattern, most phylogenetic comparative methods are not necessary for prokaryotic evolutionary studies. We are unsure how much this impression has influenced the researchers in prokaryotic genomic studies, but many papers did not use any phylogenetic comparative methods. Despite the frequent horizontal gene transfers, careful examination of the prokaryotic phylogeny could see a statistical tree (44–46). In principle, the necessity of phylogenetic comparative methods depends on the significance of the phylogenetic signal, a measure of the correlation between the evolution of the analyzed trait and the presumed phylogenetic tree. We first measured the phylogenetic signals of the analyzed traits for the 681 bacteria and 155 archaea obtained from the database TEMPURA (39). As shown in Table 1 and Additional file 1: Tables S1-S4, all the λ values are close to one, which indicates that simple statistical analysis that does not account for common ancestry’s effect would lead to inaccurate results (47, 48).

**Table 1.** The phylogenetic signals of the variables analyzed in this study.

| Traits | Bacteria |  |  | Archaea |  |  |
| --- | --- | --- | --- | --- | --- | --- |
| | *n* | Pagel's $\lambda$ | *P* | *n* | Pagel's $\lambda$ | *P* |
| Tmax | 681 | 0.957 | $3.5 \times 10^{-178}$ | 155 | 1.000 | $1.5 \times 10^{-72}$ |
| Topt, | 681 | 0.950 | $1.1 \times 10^{-196}$ | 155 | 0.988 | $5.7 \times 10^{-70}$ |
| Tmin | 681 | 0.933 | $6.6 \times 10^{-152}$ | 155 | 0.966 | $1.9 \times 10^{-53}$ |
| GC<sub>w</sub> | 681 | 1.000 | $2.6 \times 10^{-294}$ | 155 | 1.000 | $5.8 \times 10^{-60}$ |
| GC<sub>p</sub> | 681 | 1.000 | $4.7 \times 10^{-292}$ | 155 | 1.000 | $2.4 \times 10^{-59}$ |
| GC<sub>4</sub> | 681 | 1.000 | $4.8 \times 10^{-238}$ | 155 | 1.000 | $5.1 \times 10^{-53}$ |
| GC<sub>non</sub> | 681 | 1.000 | $8.6 \times 10^{-305}$ | 155 | 1.000 | $2.0 \times 10^{-65}$ |
| GC<sub>tRNA</sub> | 681 | 0.998 | $6.0 \times 10^{-275}$ | 155 | 1.000 | $2.1 \times 10^{-91}$ |
| GC<sub>5S</sub> | 646 | 1.000 | $7.0 \times 10^{-178}$ | 130 | 1.000 | $9.1 \times 10^{-51}$ |
| GC<sub>16S</sub> | 681 | 0.999 | $8.7 \times 10^{-250}$ | 155 | 0.996 | $3.7 \times 10^{-86}$ |
| GC<sub>23S</sub> | 681 | 1.000 | $2.1 \times 10^{-245}$ | 155 | 1.000 | $6.8 \times 10^{-83}$ |

Tmax, Topt, and Tmin represent maximal, optimal, and minimal growth temperature, respectively; GC<sub>w</sub>, GC<sub>p</sub>, GC<sub>4</sub>, GC<sub>tRNA</sub>, GC<sub>5S</sub>, GC<sub>16S</sub>, GC<sub>23S</sub>, and GC<sub>non</sub> represent the GC contents of the whole genome, the protein-coding sequences, the fourfold degenerate sites, the genes coding tRNAs, the genes coding 5S rRNA, the genes coding 16S rRNA, the genes coding 23S rRNA, and the non-coding DNA (including intergenic sequences and untranslated regions of mRNA), respectively. The phylogenetic signals of the chromosomal, plasmid, core and accessory genes are also very close to one and deposited in Additional file 1: Tables S1-S4.

### Bacterial but not archaeal GC contents correlated with growth temperatures

We used the phylogenetic generalized least squares (PGLS) regression to examine the relationships between GC contents and growth temperatures. The significant positive and negative slopes of the regressions correspond to significant positive and negative correlations, respectively. The slope value represents the phylogenetically corrected rate of change in GC content as growth temperature changes. Four phylogenetic models, the Brownian motion model (BM), the Ornstein-Uhlenbeck model with an ancestral state to be estimated at the root, the Pagel’s lambda model, and the early burst model, have been applied in the analysis. Their results are qualitatively identical and quantitatively similar. As the four models lead to the same conclusion, the trivial differences among their results are unrelated to understanding the relationship between GC content and growth temperature. We present the BM model results in the main text and deposit other models’ results as Additional file 1: Tables S5-S7.

Interestingly, we also found that Tmax and Topt are positively correlated with various indexes of genomic GC contents, GC_w_, GC_p_, GC_4_, and GC_non_, in bacteria (Table 2). Nevertheless, bacterial Tmin is not correlated with three GC content indexes (Table 2). In archaea, none of the three temperature indexes (Tmax, Topt, or Tmin) have any significant correlations with any of the four genomic GC content indexes (Table 2).

**Table 2.** PGLS regression of GC contents and growth temperatures.

|  | Bacteria |  |  | Archaea |  |  |
| --- | --- | --- | --- | --- | --- | --- |
| | Slope | $P$ | $P_{BH}$ | Slope | $P$ | $P_{BH}$ |
| GC<sub>w</sub>-Tmax | $7.1 \times 10^{-4}$ | $7.1 \times 10^{-4}$ | $9.4 \times 10^{-4}$ | $6.6 \times 10^{-4}$ | 0.115 | 0.153 |
| GC<sub>w</sub>-Topt | $5.7 \times 10^{-4}$ | 0.009 | 0.011 | $3.3 \times 10^{-4}$ | 0.377 | 0.503 |
| GC<sub>w</sub>-Tmin | $2.8 \times 10^{-4}$ | 0.156 | 0.226 | $5.2 \times 10^{-4}$ | 0.126 | 0.168 |
| GC<sub>p</sub>-Tmax | $6.6 \times 10^{-4}$ | 0.002 | 0.002 | $5.6 \times 10^{-4}$ | 0.183 | 0.209 |
| GC<sub>p</sub>-Topt | $5.3 \times 10^{-4}$ | 0.015 | 0.016 | $2.4 \times 10^{-4}$ | 0.522 | 0.597 |
| GC<sub>p</sub>-Tmin | $2.5 \times 10^{-4}$ | 0.202 | 0.231 | $4.6 \times 10^{-4}$ | 0.180 | 0.205 |
| GC<sub>4</sub>-Tmax | 0.001 | 0.003 | 0.003 | $9.9 \times 10^{-4}$ | 0.321 | 0.321 |
| GC<sub>4</sub>-Topt | 0.001 | 0.016 | 0.016 | $2.2 \times 10^{-4}$ | 0.806 | 0.806 |
| GC<sub>4</sub>-Tmin | $5.5 \times 10^{-4}$ | 0.239 | 0.239 | $6.9 \times 10^{-4}$ | 0.393 | 0.393 |
| GC<sub>non</sub>-Tmax | $8.0 \times 10^{-4}$ | $1.7 \times 10^{-4}$ | $2.8 \times 10^{-4}$ | $9.1 \times 10^{-4}$ | 0.025 | 0.041 |
| GC<sub>non</sub>-Topt | $6.4 \times 10^{-4}$ | 0.004 | 0.006 | $6.4 \times 10^{-4}$ | 0.080 | 0.129 |
| GC<sub>non</sub>-Tmin | $2.7 \times 10^{-4}$ | 0.170 | 0.226 | $6.5 \times 10^{-4}$ | 0.048 | 0.077 |
| GC<sub>tRNA</sub>-Tmax | $4.1 \times 10^{-4}$ | $2.2 \times 10^{-16}$ | $5.9 \times 10^{-16}$ | $7.1 \times 10^{-4}$ | $1.8 \times 10^{-11}$ | $7.2 \times 10^{-11}$ |
| GC<sub>tRNA</sub>-Topt | $3.9 \times 10^{-4}$ | $2.6 \times 10^{-14}$ | $6.9 \times 10^{-14}$ | $5.0 \times 10^{-4}$ | $2.5 \times 10^{-7}$ | $6.7 \times 10^{-7}$ |
| GC<sub>tRNA</sub>-Tmin | $1.5 \times 10^{-4}$ | $9.1 \times 10^{-4}$ | 0.002 | $4.2 \times 10^{-4}$ | $1.8 \times 10^{-6}$ | $4.7 \times 10^{-6}$ |
| GC<sub>5S</sub>-Tmax | $5.5 \times 10^{-4}$ | $1.2 \times 10^{-6}$ | $2.4 \times 10^{-6}$ | 0.001 | $1.9 \times 10^{-5}$ | $3.9 \times 10^{-5}$ |
| GC<sub>5S</sub>-Topt | $4.4 \times 10^{-4}$ | $1.4 \times 10^{-4}$ | $2.9 \times 10^{-4}$ | $8.9 \times 10^{-4}$ | $1.6 \times 10^{-4}$ | $3.2 \times 10^{-4}$ |
| GC<sub>5S</sub>-Tmin | $3.5 \times 10^{-4}$ | 0.001 | 0.002 | $6.1 \times 10^{-4}$ | 0.005 | 0.010 |
| GC<sub>16S</sub>-Tmax | $5.4 \times 10^{-4}$ | $2.2 \times 10^{-16}$ | $5.9 \times 10^{-16}$ | $8.2 \times 10^{-4}$ | $3.9 \times 10^{-11}$ | $1.0 \times 10^{-10}$ |
| GC<sub>16S</sub>-T<sub>opt</sub> | $5.2 \times 10^{-4}$ | $2.2 \times 10^{-16}$ | $8.8 \times 10^{-16}$ | $7.2 \times 10^{-4}$ | $1.1 \times 10^{-10}$ | $4.5 \times 10^{-10}$ |
| GC<sub>16S</sub>-T<sub>min</sub> | $4.6 \times 10^{-4}$ | $2.2 \times 10^{-16}$ | $8.8 \times 10^{-16}$ | $5.5 \times 10^{-4}$ | $8.5 \times 10^{-8}$ | $3.4 \times 10^{-7}$ |
| GC<sub>23S</sub>-T<sub>max</sub> | $6.6 \times 10^{-4}$ | $2.2 \times 10^{-16}$ | $5.9 \times 10^{-16}$ | 0.001 | $2.2 \times 10^{-16}$ | $1.8 \times 10^{-15}$ |
| GC<sub>23S</sub>-T<sub>opt</sub> | $6.5 \times 10^{-4}$ | $2.2 \times 10^{-16}$ | $8.8 \times 10^{-16}$ | 0.001 | $1.2 \times 10^{-14}$ | $9.5 \times 10^{-14}$ |
| GC<sub>23S</sub>-T<sub>min</sub> | $4.9 \times 10^{-4}$ | $2.2 \times 10^{-16}$ | $8.8 \times 10^{-16}$ | $8.3 \times 10^{-4}$ | $8.0 \times 10^{-11}$ | $6.4 \times 10^{-10}$ |

GC contents were the dependent variables, and growth temperatures were the independent variables.

Consistent with numerous previous studies, we found positive correlations between the GC contents of structural RNA genes (GC_tRNA_, GC_5S_, GC_16S_, and GC_23S_) and the growth temperatures (Tmax, Topt, and Tmin) in bacteria and archaea (Table 2). The significance values of these correlations are much smaller than the correlations between genomic GC content and growth temperature. Although the correlations between GC_5S_ and growth temperatures are statistically significant, their significance values are bigger than other structural RNAs. These observations indicate that the strongest correlations between GC content and growth temperatures exist in tRNAs, 16S RNA, and 23S rRNA. We noticed a rank in the slope values, from Tmax, Topt, to Tmin.

If growth temperature could shape GC contents by the stabilities of RNA secondary structures and DNA double helix, a structural RNA or a DNA double helix that is stable at the Tmax or Topt is, of course, stable at the Tmin. This logic makes it reasonable to see that the Tmin has weaker or no significant correlations with GC contents.

The difference in the correlations between bacteria and archaea might be attributed to either unknown intrinsic differences between these two domains or the substantial difference in the sample size, 681 *vs.* 155.

### Sample sizes matter

If the lack of significant correlations between genomic DNA and Tmax and Topt in archaea results from the small sample size, the correlations in bacteria would be lost when the sample size of bacteria is reduced to 155. For this reason, we randomly selected 155 bacteria from the 681 bacterial samples for 1000 rounds. The resampling analysis confirmed the idea that the sample sizes matter (Table 3; Additional file 2: Data S1). In > 950 rounds, the genomic GC content indexes (GC_w_, GC_p_, GC_4_, and GC_non_) are not correlated with Tmax or Topt (*P* > 0.05). This result could also explain the difference between the present study with Hurst and Merchant (27), which did not find significant correlations between GC_w_/GC_3_ and Topt by phylogenetic analysis of about 100 prokaryotes. Meanwhile, a few positive correlation cases happen, indicating that significant positive correlations could also be found by chance when the analyzed sample is small.

**Table 3.**
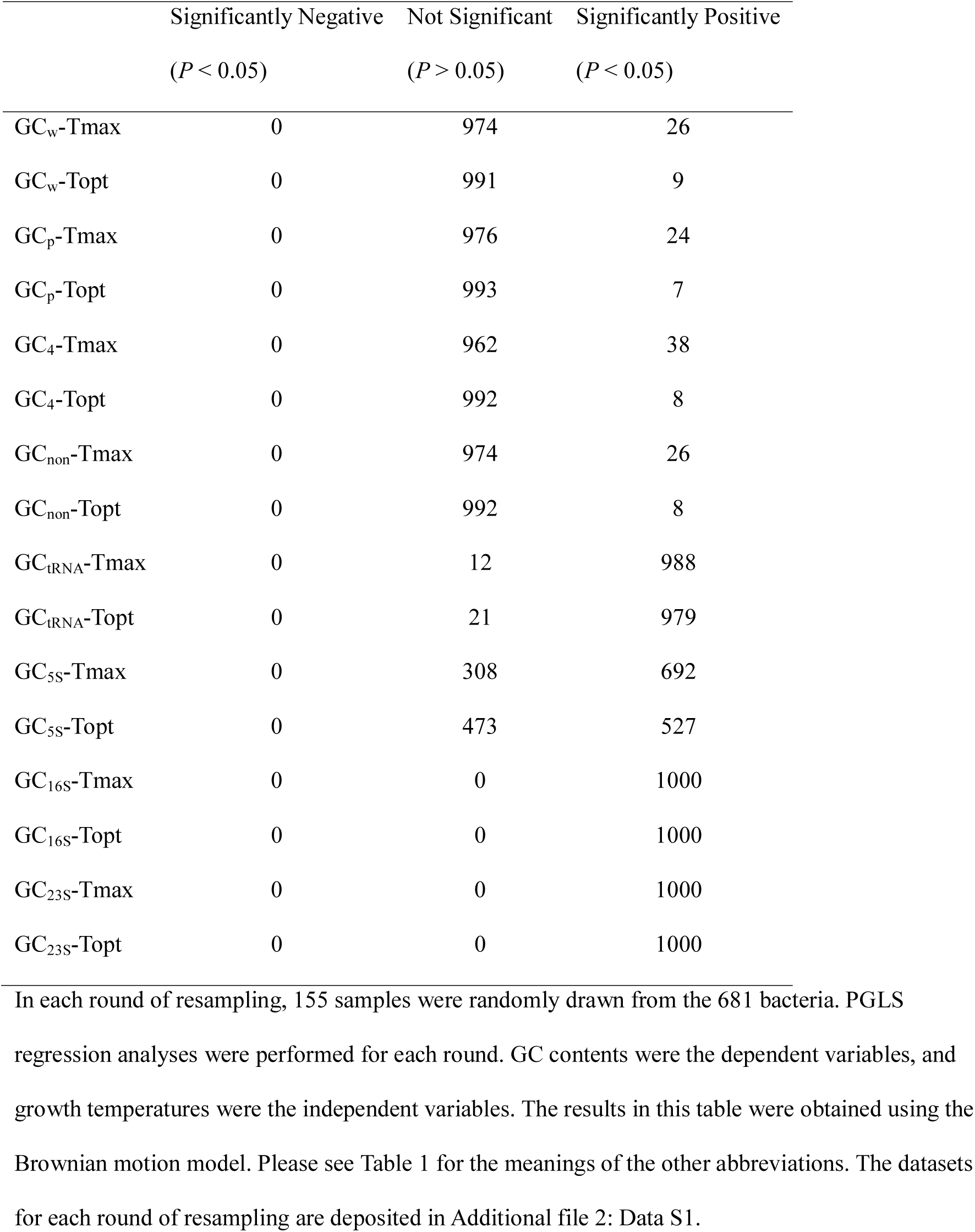
The appearance of correlations in 1000 rounds of resampling analyses.

| | Significantly Negative<br>($P < 0.05$) | Not Significant<br>($P > 0.05$) | Significantly Positive<br>($P < 0.05$) |
| --- | --- | --- | --- |
| GC<sub>w</sub>-Tmax | 0 | 974 | 26 |
| GC<sub>w</sub>-Topt | 0 | 991 | 9 |
| GC<sub>p</sub>-Tmax | 0 | 976 | 24 |
| GC<sub>p</sub>-Topt | 0 | 993 | 7 |
| GC<sub>4</sub>-Tmax | 0 | 962 | 38 |
| GC<sub>4</sub>-Topt | 0 | 992 | 8 |
| GC<sub>non</sub>-Tmax | 0 | 974 | 26 |
| GC<sub>non</sub>-Topt | 0 | 992 | 8 |
| GC<sub>tRNA</sub>-Tmax | 0 | 12 | 988 |
| GC<sub>tRNA</sub>-Topt | 0 | 21 | 979 |
| GC<sub>5S</sub>-Tmax | 0 | 308 | 692 |
| GC<sub>5S</sub>-Topt | 0 | 473 | 527 |
| GC<sub>16S</sub>-Tmax | 0 | 0 | 1000 |
| GC<sub>16S</sub>-Topt | 0 | 0 | 1000 |
| GC<sub>23S</sub>-Tmax | 0 | 0 | 1000 |
| GC<sub>23S</sub>-Topt | 0 | 0 | 1000 |

Besides, the correlations between growth temperature and the GC contents of structural RNA genes might also be lost occasionally when the sample size is severely reduced (Table 3). In the 1000 rounds of resampling, lacking significant correlations happens in 308 (for Tmax) and 473 (for Topt) rounds for 5S rRNA genes. However, in the 16S and 23S rRNA genes, positive correlations were consistently observed in all the 1000 rounds of resampling. We suspected that the tens of times more nucleotides in 16S and 23S rRNA than 5S rRNA make the results of 16S and 23S rRNAs less sensitive to small sample sizes.

In statistics, the rule of thumb boundary between small and large samples is *n* = 30. However, the results in Table 3 indicate that *n* = 155 is a too-small sample in the phylogenetic comparative analyses of the relationship between growth temperature and genomic GC content. Because of the common ancestor, two closely related lineages with similar growth temperatures and GC contents should be regarded as nearly one effective sample rather than two independent samples. The effective sample size in phylogenetic comparative studies should be much lower than the census number of the analyzed lineages.

### Positive correlations observed in genes of both chromosomes and plasmids

Previous studies showed that plasmids have significantly lower GC contents than chromosomes (8, 49, 50). Therefore, we examined the correlations between growth temperatures and GC contents separately in chromosomes and plasmids. The separations of plasmids and chromosomes are arbitrary. We strictly followed the classifications of chromosomes and plasmids of the NCBI genome database (51). Among the 681 bacteria and 155 archaea analyzed above, 172 bacteria and 42 archaea have plasmid genomes. The bacterial chromosomes also have GC contents (GC_w_, GC_p_, GC_4_, and GC_non_) positively correlated with Tmax and Topt (Table 4; Additional file 1: Tables S8-S10). Interestingly, the same pattern was also found in the bacterial plasmids (Table 4) in spite that the correlations of Tmax with GC_4_ and GC_non_ are just significant at marginal levels (0.05 < *P* < 0.1). All these correlations are not significant in archaea.

**Table 4.** PGLS regression of GC contents and growth temperatures in chromosomes and plasmids.

|  | Plasmid |  |  | Chromosome |  |  |
| --- | --- | --- | --- | --- | --- | --- |
|  | Slope | *P* | *P*<sub>BH</sub> | Slope | *P* | *P*<sub>BH</sub> |
| GC<sub>w</sub>-Tmax | 0.001 | 0.009 | 0.043 | 9.6×10<sup>-4</sup> | 0.029 | 0.043 |
| GC<sub>w</sub>-Topt | 0.001 | 0.005 | 0.031 | 9.6×10<sup>-4</sup> | 0.023 | 0.031 |
| GC<sub>p</sub>-Tmax | 0.001 | 0.016 | 0.043 | 9.1×10<sup>-4</sup> | 0.038 | 0.046 |
| GC<sub>p</sub>-Topt | 0.001 | 0.010 | 0.031 | 9.2×10<sup>-4</sup> | 0.031 | 0.034 |
| GC<sub>4</sub>-Tmax | 0.002 | 0.072 | 0.072 | 0.002 | 0.027 | 0.043 |
| GC<sub>4</sub>-Topt | 0.002 | 0.044 | 0.044 | 0.002 | 0.017 | 0.031 |
| GC<sub>non</sub>-Tmax | 8.3×10<sup>-4</sup> | 0.055 | 0.060 | 0.001 | 0.021 | 0.043 |
| GC<sub>non</sub>-Topt | 9.3×10<sup>-4</sup> | 0.025 | 0.031 | 0.001 | 0.021 | 0.031 |

GC contents were the dependent variables, and growth temperatures were the independent variables.

The common ancestor effect was not accounted for in the two previous studies comparing the GC content between plasmids and chromosomes (48, 49). By the way, we performed a phylogenetic paired t-test (52) and confirmed the pattern of lower GC content in plasmids (Additional file 1: Table S11).

### Positive correlations were observed in both core genes and accessory genes

To correspond to the previous gene-centered studies (38), we examined the correlations in bacterial core genes, i.e., genes present in all the bacteria. The number of core genes decreases rapidly with the increase in the number of analyzed bacterial genomes. With a trade-off between the number of core genes and the number of bacterial genomes, we selected 28 core genes present in 420 genomes, mostly ribosomal protein genes. Significant positive correlations have been found between GC contents (GC_p_ and GC_4_) and growth temperatures, Tmax, and Topt (Table 5; Additional file 1: Tables S12-S14).

**Table 5.** PGLS analysis of GC contents and growth temperatures in core genes and accessory genes

|  | Core Genes |  |  | Accessory Genes |  |  |
| --- | --- | --- | --- | --- | --- | --- |
|  | Slope | *P* | *P*<sub>BH</sub> | Slope | *P* | *P*<sub>BH</sub> |
| GC<sub>p</sub>-Tmax | 7.6×10<sup>-4</sup> | 9.6×10<sup>-4</sup> | 0.002 | 9.0×10<sup>-4</sup> | 0.001 | 0.002 |
| GC<sub>p</sub>-Topt | 6.4×10<sup>-4</sup> | 0.007 | 0.025 | 6.3×10<sup>-4</sup> | 0.026 | 0.030 |
| GC<sub>4</sub>-Tmax | 0.002 | 6.3×10<sup>-4</sup> | 0.002 | 0.002 | 0.003 | 0.003 |
| GC<sub>4</sub>-Topt | 0.002 | 0.004 | 0.025 | 0.002 | 0.019 | 0.030 |

GC contents were the dependent variables, and growth temperatures were the independent variables.

At the opposite side of the core genes, the accessory genes are present in one or a few bacteria. When we define the accessory genes as the genes present in less than 5% of the analyzed bacterial genomes, on average, each bacterium has 152 accessory genes. Positive correlations were observed between GC contents (GC_p_ and GC_4_) and growth temperatures (Tmax and Topt), although the values of significance are slightly larger than those in core genes (Table 5; Additional file 1: Tables S12-S14). Similar patterns were observed when we increased the threshold in defining accessory genes to 10% (*P* < 0.05 for all cases).

In addition, we compared the GC content between bacterial core genes and accessory genes using a phylogenetic paired t-test (52). Unlike the previous analysis of 36 prokaryotes that did not account for the effect of common ancestors (53), we did not observe significant differences in GC content between the core genes and the accessory genes (Additional file 1: Table S15). We also compared the chromosomal accessory genes and plasmid accessory genes. The accessory genes on chromosomes have significantly higher GC contents than those on plasmids (Additional file 1: Table S16).

### Qualitative data on growth temperature lead to the same conclusion

In the ProTraits database and the IMG database (54, 55), many prokaryotes lack quantitative measures of growth temperature but are qualitatively classified into four categories: psychrophiles/psychrotrophiles, mesophiles, thermophiles, and hyperthermophiles. We constructed a qualitative dataset of prokaryote growth temperature, including data downloaded from these two datasets and the prokaryotes in the TEMPURA database classified into the four categories referring reference (39) (Additional file 1: Table S17). We transformed the qualitative characters into numerical values by assigning 1, 2, 3, and 4 to the psychrophiles/psychrotrophiles, mesophiles, thermophiles, and hyperthermophiles, respectively. Because only some genomes have been completely assembled, we used their GC_w_ values downloaded directly from the NCBI genome database. Using the phylogenetic tree retrieved from the Genome Taxonomy Database (53), we performed PGLS regression analysis using the models mentioned above. The four models gave qualitatively identical results, so we only present the BM model because it has the smallest Akaike information criterion (AIC) value. There is a positive correlation between GC_w_ content and growth temperature in bacteria (slope = 0.457, *P* = 0.001), but not in archaea (slope = −0.582, *P* = 0.170). Although this dataset (4696 bacteria and 279 archaea) is much larger than analyzed above (681 bacteria and 155 archaea), it lost much information during the qualitative classification. All the differences in growth temperature within each category disappear.

We also examined whether the contrast in the temperature category is correlated with the contrast in the GC content between terminal tips of the phylogenetic tree. Consulting reference (6), 273 bacterial and 41 archaeal pairs were retrieved from the Genome Taxonomy Database (56). On average, the bacteria with higher ranks in Topt have 1.43% more GC than their paired bacteria with lower ranks (Additional file 1: Table S18). Pairwise comparison showed significantly higher GC contents in the bacteria with higher ranks in growth temperature (Wilcoxon signed rank test, *P* = 0.019, Fig. 1A). Still, no significant differences were observed between paired archaea with different growth temperature ranks (Wilcoxon signed rank test, *P* = 0.446, Fig. 1B).

**Figure 1.**
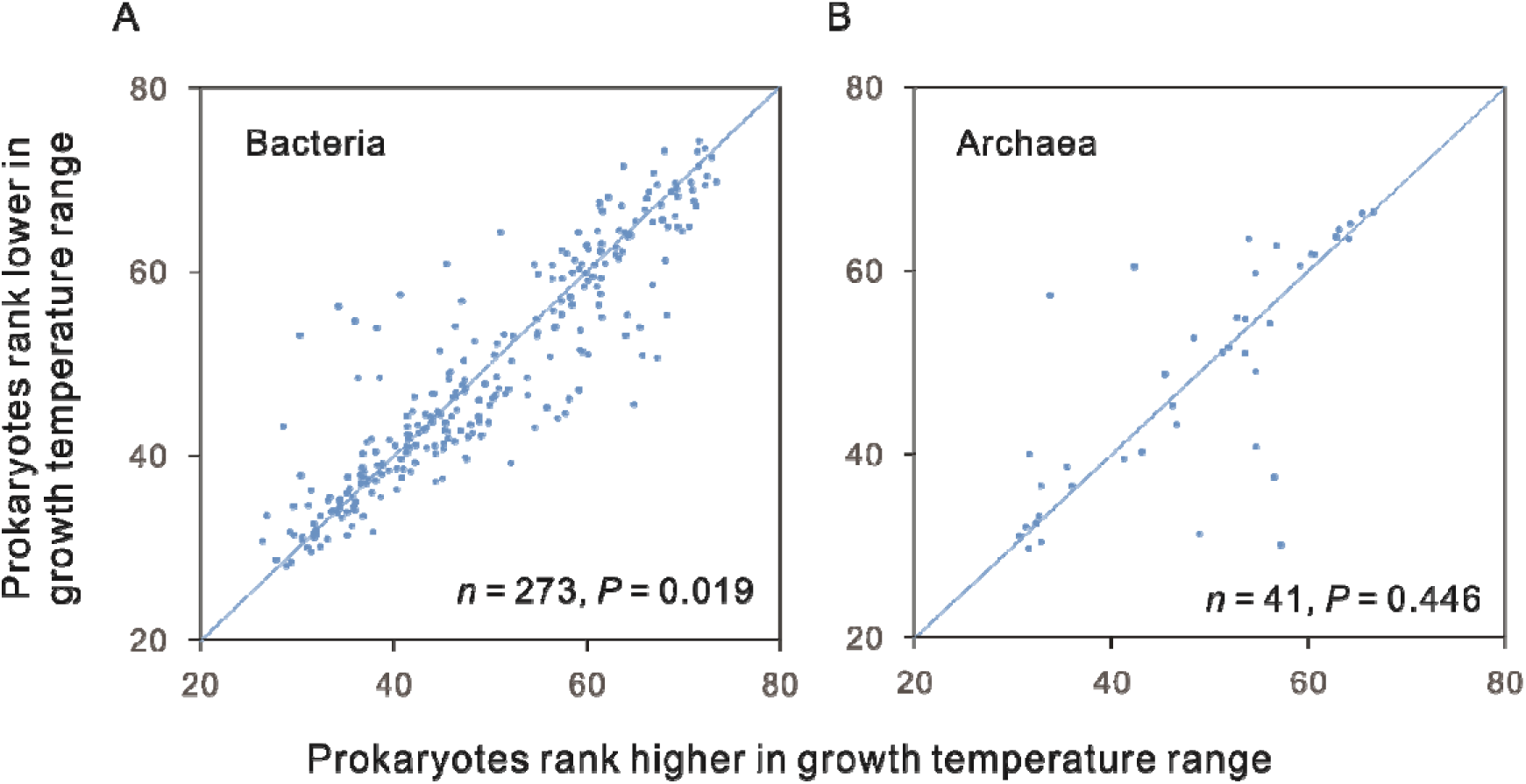
Pairwise comparison of the GC contents between closely related prokaryotes with different growth temperature ranges. Both bacteria (A) and archaea (B) were classified into four ranks according to their growth temperature, from low to high: psychrophiles/psychrotrophiles, mesophiles, thermophiles, and hyperthermophiles. The diagonal line represents cases in which prokaryotes with different ranks have the same GC contents. Points above the line (153 pairs of bacteria and 17 pairs of archaea) represent cases in which prokaryotes with higher ranks have higher GC contents than their paired relatives, while points below the line (119 pairs of bacteria and 24 pairs of archaea) indicate the reverse. The *p* values were calculated using two-tailed Wilcoxon signed-rank tests. The exact values of the GC contents are present in Additional file 1: Table S18.

### Evolutionary jumps in bacterial GC contents are correlated with Topt changes

Mahajan and Agasheand (4) recently found that the Lévy jumps model (57) could explain prokaryotic GC content evolution better than the Brownian model. The GC content constantly evolves and sometimes experiences discrete changes, i.e., jumps. Following Mahajan and Agasheand (3), we first confirmed that the Lévy jumps model could better explain the GC_w_ and Topt in our dataset than the simple Brownian model.

The Lévy jumps model has not been integrated into the PGLS packages. It could not replace the BM model in regression analysis. As an alternate, we retrieved the detected jumps in GC_w_ and examined whether significant changes in Topt accompany them. The phylogenetic locations of jumps were inferred using the *levolution* software (57). In this procedure, only the posterior probabilities (pp) of the presence of > 0 jumps were estimated, but the exact number or magnitude of jumps on each branch could not be predicted. In practice, the “precision” of jump inference is negatively correlated with the “recall” of actual jumps. By adjusting the threshold of posterior probabilities of the presence of > 0 jumps for a precision > 85%, we obtained the GC_w_ jumps with 88.5% precision and an acceptable recall of 37.0% (Additional file 1: Table S19). As shown in Fig. 2A, the magnitudes of bacterial GC_w_ jumps are positively correlated with the changes in Topt (Spearman′s rank correlation, 2-tailed, *n* = 108, rho = 0.209, *P* = 0.030). When the precision of jump inference was increased to 96.9%, the recall decreased to 21.5%, and no significant correlation was observed in the smaller sample (Spearman′s rank correlation, 2-tailed, *n* = 56, rho = 0.195, *P* = 0.150).

**Figure 2.**
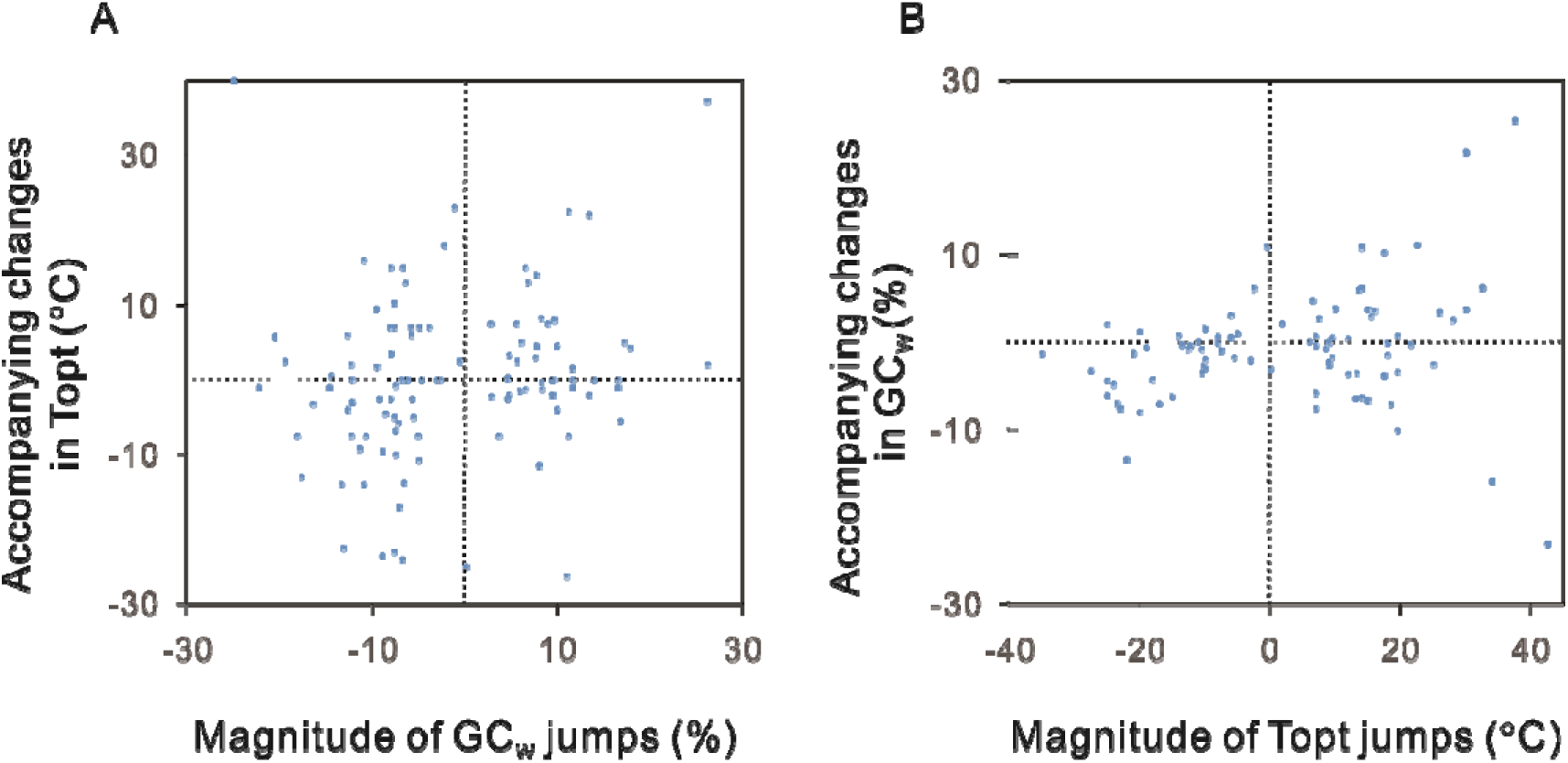
Positive correlations between the sudden changes in GC content and growth temperature of bacteria. Following Mahajan and Agasheand (3), the evolutionary jumps of GC_w_ (whole-genome GC content) and Topt (optimal growth temperature) in the bacterial phylogenetic tree were detected using the Lévy jumps model (57). (A) the magnitude of the GC_w_ jumps are significantly correlated with the accompanied changes in Topt (Spearman′s rank correlation, 2-tailed, *n* = 108, rho = 0.209, *P* = 0.030). (B) the magnitude of the Topt jumps is significantly correlated with the accompanying change in GC_w_ (Spearman’s rank correlation, 2-tailed, *n* = 86, rho = 0.280, *P* = 0.009). The exact values shown in this figure are present in Additional file 1: Tables S19-S20.

Meanwhile, we detected the evolutionary jumps in Topt using the same model. By adjusting the threshold of posterior probabilities, we inferred the Topt jumps with 95.3% precision and 21.5% recall (Additional file 1: Table S20). A positive correlation was observed between the magnitudes of the jumps in Topt and the changes of GC contents at the positions of Topt jumps (Spearman’s rank correlation, 2-tailed, *n* = 86, rho = 0.280, *P* = 0.009, Fig. 2B).

These two correlations indicate that dramatic evolutionary changes in bacterial Topt are statistically accompanied by changes in GC contents in the same direction and vice versa.

### Positive correlation between GC_w_ and Topt appears in Archaea after excluding halophiles

The above resampling analysis indicates that a positive correlation between genomic GC contents and growth temperature might be observed when we have a larger sample of archaea. In bacteria, we found such positive correlations in both chromosomes and plasmids, core and accessory genes. It seems that the positive correlation could be observed in partial sequences of bacterial genomes. For this reason, we expanded the archaeal sample size by including the incompletely assembled genomes.

For the archaea indexed in the database TEMPURA (39), we found 303 species in the All-Species Living Tree (58), a phylogenetic tree constructed using 16S rRNA sequences. These 303 samples include complete genomes and incompletely assembled genomes labeled as chromosome, scaffold, and contig in the NCBI genome database (51). First, a significant positive correlation was observed between Topt and GC_16S_ in the present dataset for all the four models we used (*P* ≤ 1.3 × 10^−6^ for all the four cases). Since many genomes of the 303 archaeal samples are not full genome sequences, we did not calculate the GC_p_, GC_4_, GC_non_, GC_tRNA_, GC_5S,_ or GC_23S_. The GC_w_ of these 303 archaea were downloaded directly from the database TEMPURA (39). Four models gave conflicting results on the relationship between GC_w_ and Topt. Only Pagel’s lambda model showed a positive correlation between GC_w_ and Topt (slope = 9 × 10^−4^, *P* = 0.02). All the other three models showed significantly negative correlations (*P* ≤ 2.2 × 10^−16^ for all the four cases). Pagel’s lambda model has the lowest AIC value and thus could be regarded as the model most fitting the data. Despite this, we are not confident in giving a conclusion based on Pagel’s lambda model.

By closely examining the scatter diagram, we noticed that Halobacteria have uniquely higher GC contents than other archaea with similar Topt (Fig. 3). The high GC content of Halobacteria was suggested to reduce the chance of thymine dimer formation caused by the intense sunlight UV irradiation (59). The strong selective force resulting from UV irradiation could overturn the potential effect of their low growth temperatures. For this reason, we examined the relationship between GC_w_ and Topt in other archaea. Although the sample size decreased to 152, the positive correlations could be observed with high confidence. Ornstein-Uhlenbeck and Pagel’s lambda models were the first and the second models most fitting the data. They all showed significant positive correlations between GC_w_ and Topt (slope = 0.001 for both cases and *P* = 0.029 and 0.046, respectively). Although the other two models, the BM model and the early burst model, did not show statistically significant correlations (*P* = 0.08 and 0.12, respectively), they presented positive slopes for the phylogenetic regressions (7.8 × 10^−4^ and 6.5 × 10^−4^).

**Figure 3.**
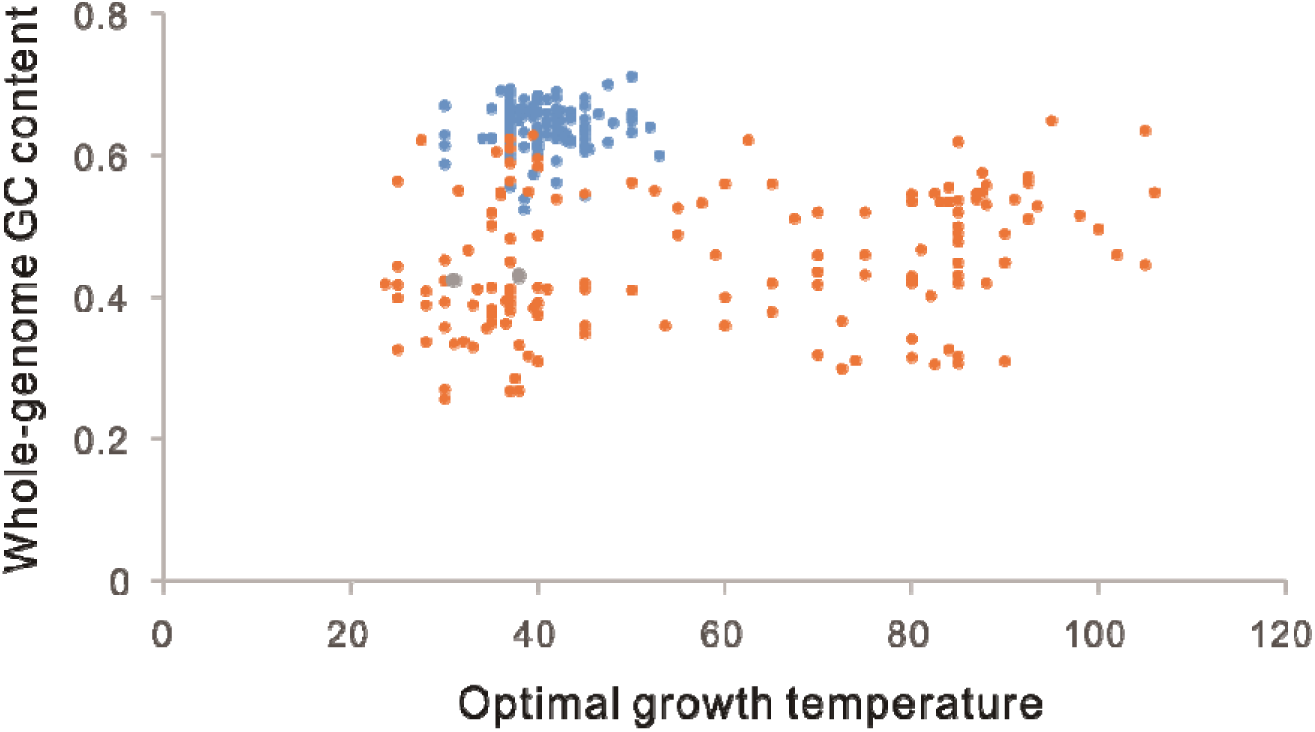
Relationship between whole-genome GC content (GC_w_) and optimal growth temperature (Topt) in Archaea. The Topt ranges of Halobacteria (*n* = 151), other halophilic archaea (*n* = 2), and nonhalophilic archaea (*n* = 150) are 30 to 53°C, 31 to 38°C, and 23.6 to 106°C, respectively. Phylogenetic generalized least squares regression analysis using the Ornstein-Uhlenbeck model with an ancestral state to be estimated at the root showed a significant positive correlation between GC_w_ and Topt in nonhalophilic archaea (slope = 0.001, *P* = 0.025).

Furthermore, we found two halophilic archaea, *Methanocalculus halotolerans* and *Methanohalophilus halophilus,* not belonging to Halobacteria, by consulting the HaloDom database (60). Excluding these two species further slightly reduced three significance values of the PGLS regression slopes, *P* = 0.025, 0.046, 0.064, and 0.10 for the Ornstein-Uhlenbeck model, the Pagel’s lambda model, the BM model (BM), and the early burst model, respectively. It should be noted that the sample size of the non-halophilic archaeal dataset is only 150. According to our resampling analysis in bacteria, it is a small size with low statistical power.

### Nonlinearity in the relationship between Topt and GC contents

PGLS regression is a method measuring linear correlations. It could just approximately show the general relationship if the correlation between GC content and Topt is nonlinear. The generalized additive mixed model (GAMM) could be used to measure the nonlinear associations across phylogenetic lineages if a low taxonomic level (e.g., species or genus) is adjusted for as a random effect (61, 62). By assuming that the species belonging to the same genus have more similar GC contents and growth temperatures than species belonging to different genera, we adjusted for the genus as a random effect. According to the genus names, the 681 bacteria and 155 archaea were divided into 536 groups. The GAMM model could give a value of the effective degrees of freedom (edf), a proxy for the degree of nonlinearity in the relationships between Topt and GC contents. An edf of 1 indicates a linear relationship, whereas a high value (8∼10 or higher) indicates high nonlinearity in the relationship (63). Using the GAMM model, we examined the nonlinear relationships between Topt and GC contents across the 836 prokaryotic genomes. As shown in Fig. 4A, the relationship between Topt and GC_w_ exhibits a moderate level of nonlinearity (edf = 5.3, *P* = 10^−4^), most likely to have some inflection points like 30□ and 70□. Similar levels of nonlinearity have been observed in the relationships of Topt with GC_p_, GC_4_, GC_tRNA_, GC_16S_, and GC_23S_ (edf = 4.5 ∼ 6.2, *P* < 0.001, Fig. 4B-F and Additional file 3: Fig. S1). The relationship between Topt and GC_5S_ exhibits a weak nonlinearity (edf = 1.5, *P* = 2 × 10^−16^). In spite of the nonlinear correlations, Topt and the GC contents of structural RNA genes exhibit clear positive associations (Fig. 4 and Additional file 3: Fig. S1). The overall trends between Topt and other GC content indexes could not be easily figured out from Fig. 4 and Fig. S1. We suggest that the above results of PGLS regressions give us the answer.

**Figure 4.**
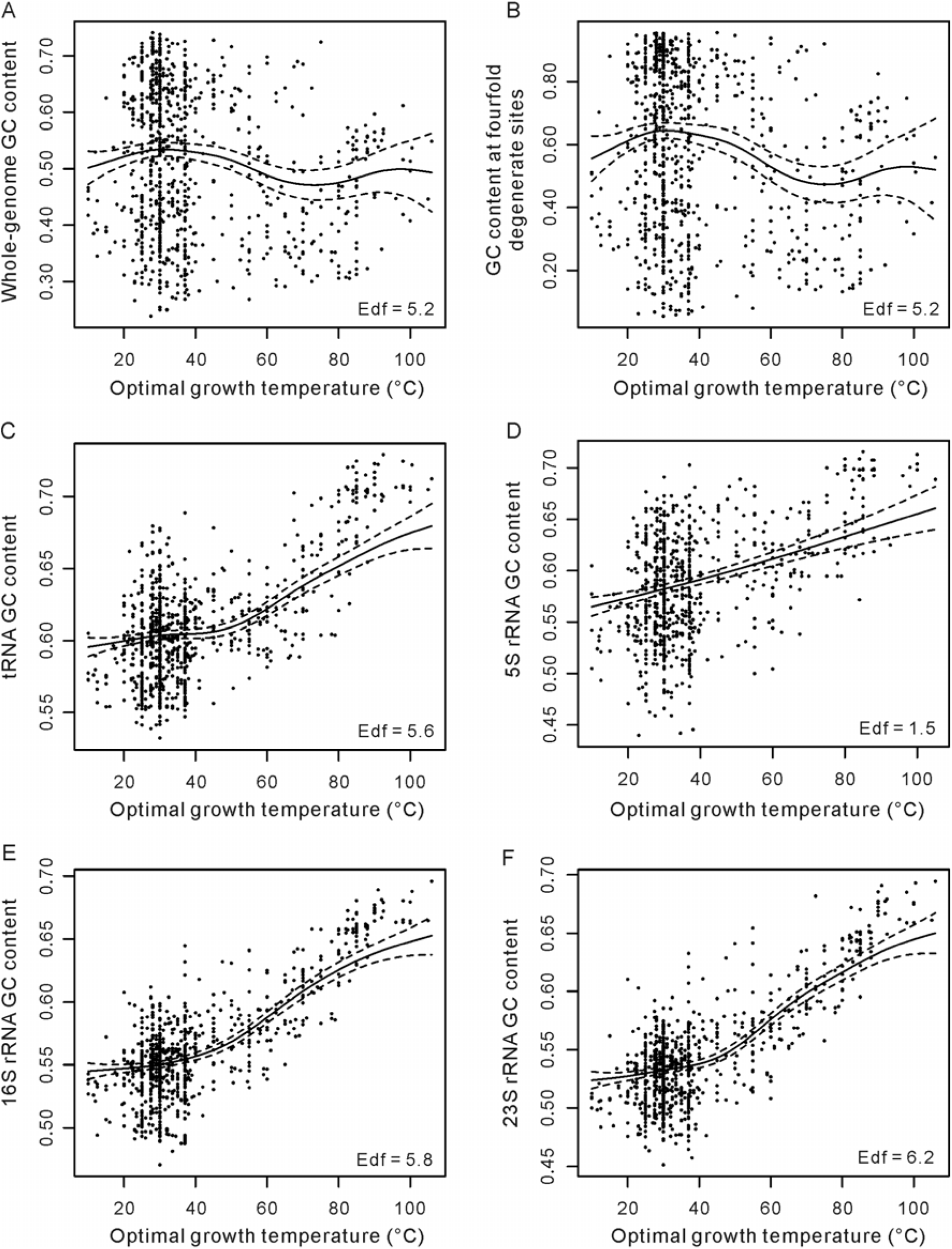
Nonlinearity in the relationship between prokaryotic optimal growth temperature and GC contents. It was estimated using the generalized additive mixed model (GAMM) by adjusting the genus as a random effect. The dataset including 836 prokaryotes (681 bacteria and 155 archaea) was used in this analysis. The 5S rRNA genes were not annotated in 60 genomes, so the analysis of the 5S rRNA has a sample size of 776. The effective degrees of freedom (edf) proxy for nonlinearity in the relationships. We presented the relationships of optimal growth temperature with the GC contents of the whole genome, fourfold degenerate sites, tRNA, 5S rRNA, 16S rRNA, and 23S rRNA as (A), (B), (C), (D), (E), and (F) in this figure and those of the protein-coding sequences and the non-coding DNA were deposited in Additional file 3: Fig. S1. The significance values of the results presented in (A) ∼ (E) are *P* = 10^−4^, 8 × 10^−7^, 2 × 10^−16^, 2 × 10^−16^, 2 × 10^−16^, and 2 × 10^−16^, respectively.

We also examined the relationships between Topt and GC contents in two subsamples of the 681 bacteria, the highest 30% Topt species and the lowest 30% Topt species using PGLS regression. No statistically significant results were obtained (*P* > 0.05 for all the cases). It may be attributed to the smaller sample sizes or the local nonlinearity of the relationship within the analyzed ranges.

### Other concerns on the correlation between GC_w_ and Topt

Some previous studies suggest that the stability of DNA double helix depends heavily on the frequency of specific dinucleotides (64–66). If GC contents influence DNA thermostability through the frequencies of specific dinucleotides, we might see positive correlations of Topt with the frequencies of some GC-content or AT-content-related dinucleotides. Referring to (67), we calculated the dinucleotide frequencies of the 681 bacterial genomes and the 155 archaeal genomes. In bacteria, no significant correlations were observed between Topt and the frequency of any dinucleotides (Additional file 1: Table S21). In archaea, only the frequency of AG(CT) is positively correlated with Topt (BM model, slope = 0.002, *P* = 0.004). This dinucleotide is not related to GC content.

We also performed multiple PGLS regression to separate archaea and bacteria as a new variable. From the dataset of 681 bacteria and 155 archaea, a scaled phylogenetic tree including 415 bacteria and 119 archaea was retrieved from TimeTree (68). GC content was the dependent variable in this regression, while the Topt and the phylogenetic domain were the two independent variables. Bacteria and Archaea were assigned to 0 and 1. PGLS regression of only two variables (GC content and Topt) was also performed as a control. Pagel’s lambda model had the lowest AIC values in both regressions, so we present the results of this model in Additional file 1: Table S22. The slope of the domain is not statistically significant (*P* > 0.220 for all cases), and the adding of this variable did not change the relationship between GC content and Topt (Additional file 1: Table S23).

Similarly, we examined whether the presence and absence of plasmid affect the relationship between GC content and Topt. As shown in (Additional file 1: Table S24), the presence and absence of plasmid is not a statistically significant variable to the evolution of GC content (*P* > 0.641 for all cases). In addition, the relationship between GC content and Topt was not affected by the presence of the second variable.

## Discussion

The GC pairs are thermally more stable than AT pairs in DNA double helix and structural RNAs (23). However, this difference is not necessarily a strong enough force to shape the evolution of GC content. As RNA structures are more sensitive to temperature elevation than DNA double helix, the growth temperature is expected to have a more substantial effect in shaping the GC content evolution of the structural RNA genes than in shaping the genomic GC content evolution. Positive correlations between growth temperature and the GC content of structural RNA genes have been repeatedly observed in various prokaryotic studies (22, 27, 39–43). However, there was a long debate on the correlation between growth temperature and genomic GC content. Benefitting from a new manual-curated dataset of prokaryotic growth temperature (39), we performed a phylogenetic comparative analysis with a much larger sample than previous studies (27, 30). In 681 bacteria, the genomic GC contents, GC_w_, GC_p_, GC_4_, and GC_non_, are all positively correlated with growth temperatures, Tmax and Topt. However, in 155 archaea, there are no significant correlations. Then, we resampled 155 bacteria from the 682 bacteria for 1000 rounds. The significant positive correlations between genomic GC contents and growth temperatures disappeared in most cases. The resampling analysis indicates that the small sample sizes of the previous analyses (27) might lead to the lack of significant correlations. It is easy to increase the sample size several times if accurate phylogenetic relationships (56) are not considered in the analysis. As shown in Table 1, we found that both growth temperatures and GC contents exhibit strong phylogenetic signals. Overlooking the effect of common ancestors would severely affect the accuracy of the results (47).

Our resampling analysis indicates that the lack of significant correlations in the 155 samples of archaea might result from the small number of effective samples. Then, we expanded the sample size to 303 archaea by including the GC contents of incompletely assembled genomes (Fig. 3). A positive correlation between GC_w_ and Topt in Archaea appears, especially after excluding the halophilic archaea. The halophilic archaea have much higher GC content than other archaea of similar growth temperature, probably because of the intense UV irradiation they have to experience (59). This result indicates that the effect of temperatures on the GC content evolution is not strong and could be easily overwhelmed by other evolutionary forces. By the same logic, we suspect that other exceptions to the positive correlation between GC content and growth temperature might have experienced some other more vital evolutionary forces shaping GC content in evolution. For example, within the hyperthermophilic genus *Thermococcus* (Topt ranging from 75 to 89°C), the GC_w_ ranges from 40.2% to 58% (39). The low-GC-content species in this genus have small genome sizes. Their low GC content might be explained by the reduced efficiency of DNA repair resulting from genome reduction and losses of DNA repairing genes (4, 69–72). In addition, GC pairs are not always more stable than AT pairs. In the presence of some ions, AT pairs become more stable than GC pairs (73). In special environmental or physiological conditions that accumulate such ions, AT-rich sequences would be more stable than GC-rich sequences, and the correlation between GC content and growth temperature would be overturned.

Besides adaptive explanations, nonadaptive processes may be explored in the future. In two ecologically distinct groups of bacteria, intracellular symbionts (including mitochondria) and marine bacterioplankton, increased AT contents are accompanied by genome reduction and gene losses (especially the losses of DNA repair genes like *mutY* genes) (4, 69–72). If most DNA damages tend to decrease GC content, as suggested by some previous studies (74, 75), many DNA repairing genes would counter such effects or even increase GC content as documented in gene conversions (10, 17). Heat stress can lead to various DNA damages from deaminated cytosine, 8-oxoguanine, to single- and double-stranded DNA breaks (76, 77). Together with the sensitivity of macromolecular stability to increased temperature, thermophiles experienced a strong selective force for low mutation rates (78, 79) and efficient DNA repair systems. We propose that an increase of DNA repair efficiency associated with the increasing growth temperature, or a decrease of DNA repair efficiency associated with the decreasing growth temperature, might shape the evolution of GC content evolution like that happens in intracellular symbionts and marine bacterioplankton.

A recent study suggests that sequential amino acid substitutions are involved in the thermal adaptation in the archaeal order *Methanococcales* and revealed arginine as the most favored amino acid (80). As six GC-rich codons encode the arginine, the thermal adaptation at the proteomic level would affect the evolution of genomic GC content. Because the 4-fold degenerate sites are free from the evolutionary forces coming from the natural selection acting on protein sequences, our observations of similar correlations of GC_w_, GC_p_, and GC_4_ with growth temperature indicate that the nucleotide composition evolved independently in bacterial adaptation to high temperatures.

As the frequent gain and loss of plasmids, the plasmid DNAs could be regarded as accessory genomes. Because of the high turnover rates of plasmids and accessory genes in prokaryotic evolution, we could regard them as new immigrants, as opposed to the natives for the chromosomes and core genes. Although the core genes and even the ribosomal RNA genes may occasionally be transferred across different prokaryotic lineages (81, 82), the fitness cost of inter-species replacement of homologous sequences (83) restricts the frequency of the core genes. Genes performing essential informational tasks in the cell are less frequently transferred across lineages (84, 85). Our phylogenetic correlation analysis showed positive correlations between GC contents and growth temperatures in chromosomes, core genes, plasmids, and accessory genes. Also, there is no sharp difference in the correlations between the new immigrants and the natives. In large-scale analyses of horizontal gene transfer in prokaryotes, GC-content similarity between donor and recipient was found to be the factor, or one of the factors, governing the compatibility of the new immigrants in new hosts (86, 87). The effect of promoter GC content on the expression of the new immigrants was suggested to be the underlying mechanism governing the compatibility (88). Here, we suggest that the temperature-associated structural stabilities, including the stability of DNA double helix, the stability of the transient DNA-RNA duplex during transcription, and maybe the stability of the possible secondary structures of mature mRNA (1), might be another nonexclusive factor governing the compatibility. The new immigrants compatible with the host should have GC contents adapted to the host’s growth temperature.

A previous serial transfer experiment seems to be contradictory to our results. Increased genomic GC content was not observed in the bacterium *P. multocida* after 14,400 generations of increasing temperature from 37°C to 45°C (28). Although we observed a positive correlation between genomic GC content and growth temperature, we do not think a small increment in GC content, resulting from either a GC-biased mutator or integration of a GC-rich exogenous sequence, would bring a great advantage to the host organism. Most likely, it is just a slight advantage. According to the population genetic theory, the slightly beneficial mutants are efficiently selected only when they are in a large population. The experimental evolution generally involves severe, periodic reductions in population size, and the bottleneck effect dramatically reduces the fixation probability of beneficial mutations (89). As we see, large-scale statistical analysis has the advantage of revealing slightly beneficial traits.

Musto et al. (30) emphasize that the growth temperature can be the only influencing factor in GC content evolution only when closely related species are compared. Our pairwise comparison of neighboring branches with different ranks of growth temperature (Fig. 1) gave the same conclusion as our PGLS analyses. We agree that many factors influence GC content evolution, and the positive relationship between growth temperature and GC content is statistically significant. In the 273 pairs of bacteria, there are 153 pairs where high growth temperature ranks have higher GC contents and 119 pairs with the opposite pattern.

Mahajan and Agasheand (3) and the present study found that the evolutionary rates of GC content and growth temperature have occasional jumps assumed in the Lévy jumps model (57). As shown in Fig. 2, the jump-ups and jump-downs of GC content are significantly correlated with changes in growth temperature and vice versa. It should be emphasized that not all increases in growth temperature were accompanied by increases in GC content. There are just statistically significant correlations (*P* < 0.05).

Some dinucleotides could significantly enhance the stability of double-strand DNA (64–66). To examine whether the effect of some dinucleotides underlies the positive correlation between GC content and growth temperature, we examined the relationship between Topt and the frequencies of dinucleotides. Unfortunately, no GC-content-altering dinucleotides meet the expectation.

## Conclusions

We should remark that what we observed are weak correlations between genomic GC content and growth temperature. The slopes of the PGLS regressions are generally between 10^−3^ and 10^−4^. The bacteria rank higher in growth temperature have just 1.43% more GC (Fig. 1A). Considering the significant difference in the thermoresistence of nucleic acids between *in vivo* and *in vitro* (36), we believe that other cellular components mainly contribute to the thermostability of nucleic acids in thermophiles and hyperthermophiles, and the increase of GC content just plays a supplemental role. Moreover, we observed correlations between GC content and growth temperature, suggesting rather than proving the causal effects between the two variables. We should be open to the thermal adaptation hypothesis (23) and other intricate explanations, including nonadaptive ones. This paper aims to end the long-standing debate on the relationship between GC content and growth temperature. Only after establishing the positive correlation could the attention of genome biologists be paid to the biological significance of the correlation.

## Methods

We downloaded the prokaryote growth temperatures from the database TEMPURA (39). This database contains 8,639 manual curated prokaryotes (549 archaea and 8090 bacteria). Using the links to the NCBI Taxonomy database (90) and the taxonomy IDs provided by TEMPURA for each prokaryotic strain, we obtained 1110 prokaryotes whose genome assembly levels were labeled as“complete” from the NCBI database (91). Among them, we found the phylogenetic information for 682 bacteria and 156 archaea from Genome Taxonomy Database (56). The sequences of these genomes were downloaded from the NCBI genome database (51). To avoid annotation bias resulting from different methods, all the genomes were re-annotated using the DFAST, version 1.2.11, with its default parameters (92). In total, we obtained the genome annotations for 836 prokaryotes (681 bacteria and 155 archaea). The GC contents of these prokaryotes were calculated from their genome sequences. The genomes accession numbers and the database links for the 836 prokaryotes are deposited in Additional file 1: Table S25.

We also constructed a large dataset according to their growth temperature qualitatively. First, we divided the 836 prokaryotes (from the database TEMPURA) mentioned above into four categories according to their growth temperature referring to (39): psychrophiles/psychrotrophiles (Topt < 20°C), mesophiles (20 ≤ Topt < 45°C), thermophiles (45 ≤ Topt < 80°C), and hyperthermophiles (80°C ≤ Topt). Then, we downloaded the lists of prokaryotes labeled with psychrophiles/psychrotrophiles, mesophiles, thermophiles, or hyperthermophiles from the ProTraits database and the IMG database (54, 55). Then, we combined the datasets from these three sources (TEMPURA, ProTraits, and IMG) and discarded the overlapping items, the conflicting items, and the items lacking phylogenetic information in the Genome Taxonomy Database (56). Finally, we obtained a new dataset including 4696 bacteria and 279 archaea (Additional file 1: Table S17). The whole-genome GC contents of these prokaryotes were downloaded directly from the genome report file of the NCBI genome database (93)

As the contrasts between different pairs of terminal tips of the phylogenetic tree are independent, pairwise comparisons between pairs of terminal tips could control the effect of common ancestors. Referring to reference (6), we wrote a script to select pairs of closely related bacteria with different ranks of growth temperature (psychrophiles/psychrotrophiles, mesophiles, thermophiles, and hyperthermophiles). In cases where two or more neighboring tips with the same rank were used to pair with bacteria with another rank, we used the average value of their GC contents to represent the GC content of their internal node. The script is deposited as Additional file 4: Data S2.

The phylogenetic signals (λ) of both GC contents and growth temperatures were estimated using the *phylosig* function of the R (Version 4.0.3) package *phytools* (Version 0.7-70) (94). The PGLS regression was performed using the R (Version 4.0.3) package *phylolm* (version 2.6.2) with the default parameters (95).

To avoid false-positive results that might happen in multiple correlation analyses of the same dataset, we controlled the false discovery rate by the Benjamini-Hochberg (BH) procedure using the p.adjust function in R (Version 4.0.3).

Following Mahajan and Agasheand (3), we used the *geiger* package (96) and the *levolution* software (57) to simulate our datasets, estimate the branch-specific posterior probabilities of jumps and infer the phylogenetic location of jumps.

The GAMM regressions were performed using the *gamm4* function of the package *gamm4* (Version 0.2-6, based on package *mgcv* and package *lme4*). The formula is:

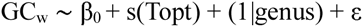

where the GC_w_ is the response variable, and the Topt is the explanatory variable, s(X) means that a smoothing function is used for the explanatory variable. The expression (1|genus) means that a random component was specified with genus as random effects.

## Abbreviations

A: adenine
BM: Brownian motion model
C: cytosine
G: guanine
GC_16S_: GC contents of the genes coding 16S rRNA
GC_23S_: GC contents of the genes coding 23S rRNA
GC_3_: GC content at the third site of the codons
GC_4_: GC contents of the fourfold degenerate sites
GC_5S_: GC contents of the genes coding 5S rRNA
GC_non_: GC contents of the non-coding DNA (including intergenic sequences and untranslated regions of mRNA)
GC_p_: GC contents of the protein-coding sequences
GC_tRNA_: GC contents of the genes coding tRNAs
GC_w_: GC contents of the whole genome
PGLS: phylogenetic generalized least squares
T: thymine
Tmax: maximal growth temperature
Tmin: minimal growth temperature
Topt: optimal growth temperature

## Declarations

### Ethics approval and consent to participate

Not applicable.

### Consent for publication

Not applicable.

### Availability of data and materials

All data generated or analyzed during this study are included in this published article and its supplementary information files.

### Authors’ contributions

DKN conceived the study and wrote the manuscript. EZH, XRL, ZLL, and JG performed the data analysis. All authors read, improved, and approved the final manuscript.

## Acknowledgements

Not applicable.

