## Supplementary Data S1 for "A positive correlation between GC content and growth temperature in prokaryotes": Readme.txt

We randomly selected 155 bacteria from the 681 bacterial samples for 1000 rounds by using the sample function in R (Version 4.0.3).
Then，we used the phylogenetic generalized least squares (PGLS) regression to examine the relationships between GC contents and growth temperatures in the 1000 rounds.Brownian motion model (BM) and the Pagel's lambda model (lambda) have been applied in this analysis.
In the table, temp_tmax, temp_toa, and temp_tmin represent maximal, optimal, and minimal growth temperature, respectively; genome_gc_content, mrna_gc_content, ffold_gc_content, trna_gc_content, rrna5s_gc_content, rrna16s_gc_content, rrna23s_gc_content, and noncod_gc_content represent the GC contents of the genome, the protein-coding sequences, the fourfold degenerate sites, the genes coding tRNAs, the genes coding 5S rRNA, the genes coding 16S rRNA, the genes coding 23S rRNA, and the non-coding DNA (including intergenic sequences and untranslated regions of mRNA), respectively.
