## supplementary figure S1 for "A positive correlation between GC content and growth temperature in prokaryotes"

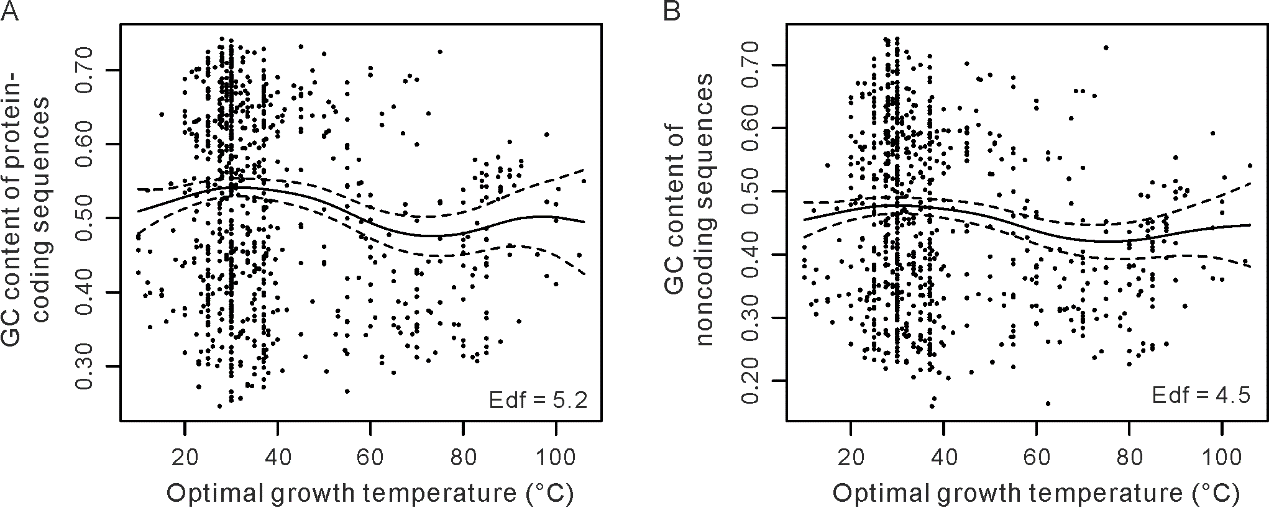

Fig. S1. Nonlinearity in the relationship between prokaryotic optimal growth temperature and GC contents. It was estimated using the generalized additive mixed model (GAMM) by adjusting for the genus as a random effect. The dataset including 681 bacterial and 155 archaeal species was used in this analysis. The effective degrees of freedom (edf) is a proxy for the level of nonlinearity in the relationships. We presented the relationships of optimal growth temperature with the GC contents of protein-coding sequences and non-coding DNA (intergenic sequences and untranslated regions of mRNA that are generally unannotated in prokaryotic genomes) as (A) and (B) in this figure. The significance values of the results presented in (A) and (B) are *P* = 5 × 10^−5^ and 9 × 10^−4^, respectively.
