## Supplementary Data S2 for "A positive correlation between GC content and growth temperature in prokaryotes": find_pgls_pair_by_depth_final.py

```
##This script is used to analyze the correlation between traits.
##The first command-line argument is the stardard trait, which must be one of all trails.
##The second command-line argument is the file of the species trait information, first column is the species name, and the other columns are the trait. The columns are separated by tab, and the presence or absence of a trait is represented by 0/1, and numerical trait is represented by numbers.
##The third command-line argument is the file of the phylogenetic tree, which needs to be in NWK format.
##The fourth command-line argument is the number of cycles to be searched, which is represented by numbers
##for example:
##python3 find_pgls_pair_by_depth_final.py group bac_tempura_pair.csv bac_tempura_pair.nwk 2
from Bio import Phylo
import copy
import sys
import scipy.stats

def get_parent(TREE,CLADE):
	path_nodes = TREE.get_path(CLADE)
	return path_nodes[-2]

def is_brother(TREE,CLADE1,CLADE2):
	return get_parent(TREE,CLADE1) == get_parent(TREE,CLADE2)

def check_pair(CLADE_A,CLADE_B,DATA,STANDARD):
	tags = []
	clades_list = [CLADE_A,CLADE_B]
	for clade in clades_list:
		clade_name = clade.name
		tag = DATA[clade_name][STANDARD]
		tags.append(tag)
	tags_set = set(tags)
	return len(tags_set) > 1

def read_list(FILE):
	DATA = {}
	fo = open(FILE,"r")
	for line in fo :
		line = line.strip()
		if line.startswith("name"):
			CULUMNS = line.split(',')
		else:
			species_name = line.split(',')[0]
			DATA[species_name] = {}
			for para in CULUMNS[1:]:
				try :
					DATA[species_name][para] = float(line.split(',')[CULUMNS.index(para)])
				except:
					DATA[species_name][para] = str(line.split(',')[CULUMNS.index(para)])
	fo.close()
	return(DATA,CULUMNS)

def search_by_level(TREE,DATA,STANDARD,OUT):
	out = open(OUT,"a")
	all_ter_clades = list(TREE.find_clades(terminal = True,order = "level"))
	lookup = {}
	for i,elem in enumerate(all_ter_clades):
		lookup[elem] = i
	uplook = {value: key for key, value in lookup.items()}
	prune_list = []	
	pgls_list = []
	for ter_clade_a in all_ter_clades:
		if len(TREE.get_path(ter_clade_a)) > 1:
			for ter_clade_b in all_ter_clades[lookup[ter_clade_a]:]:
				if ter_clade_a != ter_clade_b and is_brother(TREE,ter_clade_a,ter_clade_b):
					if check_pair(ter_clade_a,ter_clade_b,DATA,STANDARD):
						pgls_list.append((ter_clade_a,ter_clade_b))
					else:
						prune_list.append((ter_clade_a,ter_clade_b))
					break
	for clade_c,clade_d in pgls_list:
		if DATA[clade_c.name][STANDARD] == 1:
			out.write(clade_c.name + "\t" + clade_d.name)
			out.write("\t")
			for trait in  DATA[clade_c.name].keys():
				out.write(str(DATA[clade_c.name][trait]) + "\t" + str(DATA[clade_d.name][trait]))
				out.write("\t")
#			out.write(str(TREE.distance(clade_c,clade_d)))
			out.write("\n")
			

		else:
			out.write(clade_d.name + "\t" + clade_c.name)
			out.write("\t")
			for trait in DATA[clade_c.name].keys():
				out.write(str(DATA[clade_d.name][trait]) + "\t" + str(DATA[clade_c.name][trait]))
				out.write("\t")
#			out.write(str(TREE.distance(clade_d,clade_c)))
			out.write("\n")
		TREE.prune(clade_c)
		TREE.prune(clade_d)

	for clade_e,clade_f in prune_list:
		old_name_e = clade_e.name
		old_name_f = clade_f.name
		new_name = old_name_e + "+" + old_name_f
		DATA[new_name] = {}
		for trait in DATA[clade_e.name].keys():
			if  isinstance(DATA[clade_e.name][trait],float):
				new_value = round((DATA[clade_e.name][trait] + DATA[clade_f.name][trait])/2,5)
				DATA[new_name][trait] = new_value
			else:
				new_value = DATA[clade_e.name][trait] + "+" + DATA[clade_f.name][trait]
				DATA[new_name][trait] = new_value

		TREE.prune(clade_e)
		clade_f.name = new_name
	out.close()
	return(TREE,DATA,STANDARD)

in_file_name = sys.argv[2]
standard = sys.argv[1]
tree_name = sys.argv[3]
original_tree = Phylo.read(tree_name,"newick")
repeat_num = int(sys.argv[4])
tree = copy.deepcopy(original_tree)
data,culumns = read_list(in_file_name)
out_file_name = "./search_result.txt"
out = open(out_file_name,"w")
out.write("#")
for cul in culumns:
	out.write(cul)
	out.write("\t")
out.write("\n")
out.close()
for i in range(repeat_num):
	tree,data,standard = search_by_level(tree,data,standard,out_file_name)
```
